# Updated vaccine protects from infection with SARS-CoV-2 variants, prevents transmission and is immunogenic against Omicron in hamsters

**DOI:** 10.1101/2021.11.12.468374

**Authors:** Sapna Sharma, Thomas Vercruysse, Lorena Sanchez-Felipe, Winnie Kerstens, Madina Rasulova, Rana Abdelnabi, Caroline S Foo, Viktor Lemmens, Dominique Van Looveren, Piet Maes, Guy Baele, Birgit Weynand, Philippe Lemey, Johan Neyts, Hendrik Jan Thibaut, Kai Dallmeier

**Affiliations:** KU Leuven Department of Microbiology, Immunology and Transplantation, Rega Institute, Laboratory of Virology, Molecular Vaccinology and Vaccine Discovery, Leuven, Belgium; KU Leuven Department of Microbiology, Immunology and Transplantation, Rega Institute, Laboratory of Virology and Chemotherapy, Translational Platform Virology and Chemotherapy, Leuven, Belgium; KU Leuven Department of Microbiology, Immunology and Transplantation, Rega Institute, Laboratory of Clinical and Epidemiological Virology, Zoonotic Infectious Diseases Unit, Leuven, Belgium; KU Leuven Department of Microbiology, Immunology and Transplantation, Rega Institute, Laboratory of Clinical and Epidemiological Virology, Evolutionary and Computational Virology, Leuven, Belgium; KU Leuven Department of Imaging and Pathology, Translational Cell and Tissue Research, B-3000 Leuven, Belgium; Global Virus Network (GVN), Baltimore, MD, USA

**Keywords:** SARS-CoV-2, variants of concern (VOC), vaccine efficacy, antigenic cartography, virus transmission, hamster model, second-generation vaccine

## Abstract

Current first-generation COVID-19 vaccines are based on prototypic spike sequences from ancestral 2019 SARS-CoV-2 strains. However, the ongoing pandemic is fueled by variants of concern (VOC) that threaten to escape vaccine-mediated protection. Here we show in a stringent hamster model that immunization using prototypic spike expressed from a potent YF17D viral vector (1) provides vigorous protection against infection with ancestral virus (B lineage) and VOC Alpha (B.1.1.7), however, is insufficient to provide maximum protection against the Beta (B.1.351) variant. To improve vaccine efficacy, we created a revised vaccine candidate that carries an evolved spike antigen. Vaccination of hamsters with this updated vaccine candidate provides full protection against intranasal challenge with all four VOCs Alpha, Beta, Gamma (P.1) and Delta (B.1.617.2) resulting in complete elimination of infectious virus from the lungs and a marked improvement in lung pathology. Vaccinated hamsters did also no longer transmit the Delta variant to non-vaccinated sentinels. Hamsters immunized with our modified vaccine candidate also mounted marked neutralizing antibody responses against the recently emerged Omicron (B.1.1.529) variant, whereas the old vaccine employing prototypic spike failed to induce immunity to this antigenically distant virus. Overall, our data indicate that current first-generation COVID-19 vaccines need to be urgently updated to cover newly emerging VOCs to maintain vaccine efficacy and to impede virus spread at the community level.

**Significance Statement:** SARS-CoV-2 keeps mutating rapidly, and the ongoing COVID-19 pandemic is fueled by new variants escaping immunity induced by current first-generation vaccines. There is hence an urgent need for universal vaccines that cover variants of concern (VOC). In this paper we show that an adapted version of our vaccine candidate YF-S0* provides full protection from infection, virus transmission and disease by VOCs Alpha, Beta, Gamma and Delta, and also results in markedly increased levels of neutralizing antibodies against recently emerged Omicron VOC in a stringent hamster model. Our findings underline the necessity to update COVID-19 vaccines to curb the pandemic, providing experimental proof on how to maintain vaccine efficacy in view of an evolving SARS-CoV-2 diversity.

## Introduction

Severe Acute Respiratory Syndrome Corona Virus 2 (SARS-CoV-2) emerged as a zoonosis likely from a limited number of spill-over events into the human population (2). Nevertheless, the ongoing COVID-19 pandemic is entirely driven by variants that evolved during subsequent large-scale human-to-human transmission. In particular, mutations within the viral spike protein are under continuous surveillance considering their role in viral pathogenesis and as target for virus-neutralizing antibodies (nAb). Following early diversification, the D614G SARS-CoV-2 variant (B.1 lineage) became dominant in March 2020. Late 2020, Variants of Concern (VOC) emerged with increased transmissibility, potentially increased virulence and evidence for escape from naturally acquired and vaccine-induced immunity (3). This involved four VOCs harboring a unique set of partially convergent, partially unique spike mutations as compared to prototypic (Wuhan) or early European D614G (B.1) lineages of SARS-CoV-2, namely VOCs Alpha (B.1.1.7; N501Y D614G), Beta (B.1.351; K417N E484K N501Y D614G), Gamma (P.1; K417T E484K N501Y D614G) and Delta (B.1.617.2; K417T L452R T478K D614G P681R)(4). N501Y was first detected in VOC Alpha and has been linked to an enhanced transmissibility due to an increased affinity for the human ACE-2 receptor (5, 6). Subsequent emergence of E484K within this lineage hampers the activity of nAb suggestive for immune escape (7-9). Likewise, a combination of K417N and E484K (10) may explain a marked reduction in vaccine efficacy (VE) of some vaccines such as ChAdOx1 nCoV-19 (AstraZeneca, Vaxzevria) in clinical trials in South Africa during high prevalence of VOC Beta (11). Similarly, sera from vaccinees immunized with first-generation mRNA (Pfizer-BioNTech, Cormirnaty; Moderna, mRNA-1273) or nanoparticle subunit vaccines (Novavax) showed a substantial drop in neutralizing capacity for VOC Beta (12). Furthermore, VOC Gamma harboring K417T and E484K emerged in regions of Brazil with high seroprevalence, so despite naturally acquired immunity against prototypic SARS-CoV-2 (13). VOC Delta was first identified in October 2020 in India (14) and became the predominant SARS-CoV-2 lineage worldwide in 2021, driven by a substantially increased transmissibility (15). In late November 2021, a new VOC Omicron (B.1.1.529) was discovered in southern Africa (16) and is since then spreading globally; displacing other strains at unprecedented speed. Omicron carries by far the largest number (>32) of mutations, deletions, and insertions in its spike protein described to date (17, 18), including a combination of substitutions previously linked to increased human-to-human transmission (N501Y D614G P681H) as well as escape from antibody-mediated immunity (K417N E484A) acquired by natural exposure or elicited current vaccines (19, 20). While the intrinsic pathogenic potential of Omicron remains uncertain (21), its antigenic divergence leads to a loss of activity of most therapeutic monoclonal antibodies (22) and failure of current first-generation vaccines to protect from infection (23, 24). The maintenance of some cross-protective nAb levels may require repeated booster dosing (24-26).

Currently licensed COVID-19 vaccines and vaccine candidates in advanced clinical development are based on antigen sequences of early SARS-CoV-2 isolates that emerged in 2019 (27). We reported on a YF17D-vectored SARS-CoV-2 vaccine candidate using prototypic spike as vaccine antigen (YF-S0; S0) that had an outstanding preclinical efficacy against homologous challenge (1). However, in the current study we demonstrate to what extent VE of S0, and hence first-generation spike vaccines in general, may decline when trialed against VOC in a stringent hamster model (28). Therefore, we designed a second-generation vaccine candidate (YF-S0*) by (*i*) modifying its antigen sequence to keep in pace with the evolving spike variant spectrum, in combination with (*ii*) a further stabilized protein conformation (29). This new S0* vaccine candidate provides full protection against all current VOCs Alpha, Beta, Gamma and Delta. Since Omicron does not cause a productive infection nor apparent pathology in hamsters (30), levels of nAb which increased markedly after vaccination with S0* serve as proxy for an improved VE. Finally, hamsters vaccinated with S0* do no longer transmit the virus to non-vaccinated sentinels during close contact, even under conditions of prolonged co-housing and exposure to a high infectious dose of VOC Delta.

It is a challenging task to develop a universal vaccine that follows the evolution of emerging SARS-CoV-2 variants such as Omicron that carry increasingly complex combinations of old and new driver mutations responsible for both nAb escape (e.g., E484K/A) (10) and enhanced transmission (e.g., N501Y; P681R/H) (31). However, our findings provide strong experimental support that first-generation COVID-19 vaccines need to be urgently adapted for VE against current and future VOCs fueling the ongoing pandemic.

## Results and Discussion

### No change regarding VOC Alpha, yet markedly reduced efficacy of first-generation spike vaccine against VOC Beta

To assess VE of prototypic spike antigen against VOCs, hamsters were vaccinated twice with each 10^4^ PFU of YF-S0 (S0) or sham at day 0 and 7 via the intraperitoneal route (1) (**Fig. 1A**). Serological analysis at day 21 confirmed that 30/32 (94%) vaccinated hamsters had seroconverted to high levels of nAbs against prototypic SARS-CoV-2 with geometric mean titre (GMT) of 2.3 log_10_ (95% CI 2.0-2.6) (**Fig. 1B**). Next, animals were challenged intranasally with 1×10^3^ TCID_50_ of either prototypic SARS-CoV-2, VOC Alpha or Beta as established and characterized before in the hamster model (28). At day four after infection (4 dpi), viral replication was determined in lung tissue by qPCR and virus titration (**Fig. 1C, D**). In line with what was originally described for S0 (1), a marked reduction in viral RNA and infectious virus loads down to undetectable levels (up to 6 log_10_ reduction) was observed in the majority of animals challenged with either prototypic SARS-CoV-2 (8/10; 86% VE) or VOC Alpha (9/10; 88% VE). In those animals (2/10 and 1/10, respectively) that were not completely protected, virus loads were at least 100 times lower than in infected sham controls. By contrast and despite full immunization, S0 vaccination proved to be less effective against VOC Beta, with only 4/12 hamsters without detectable infectious virus (60% VE). Nonetheless, in the remaining 8/12 animals with breakthrough infection by VOC Beta, viral replication was tempered as vaccination still resulted in a 10 to 100-fold reduction in infectious virus titres relative to sham.

**Figure 1.**
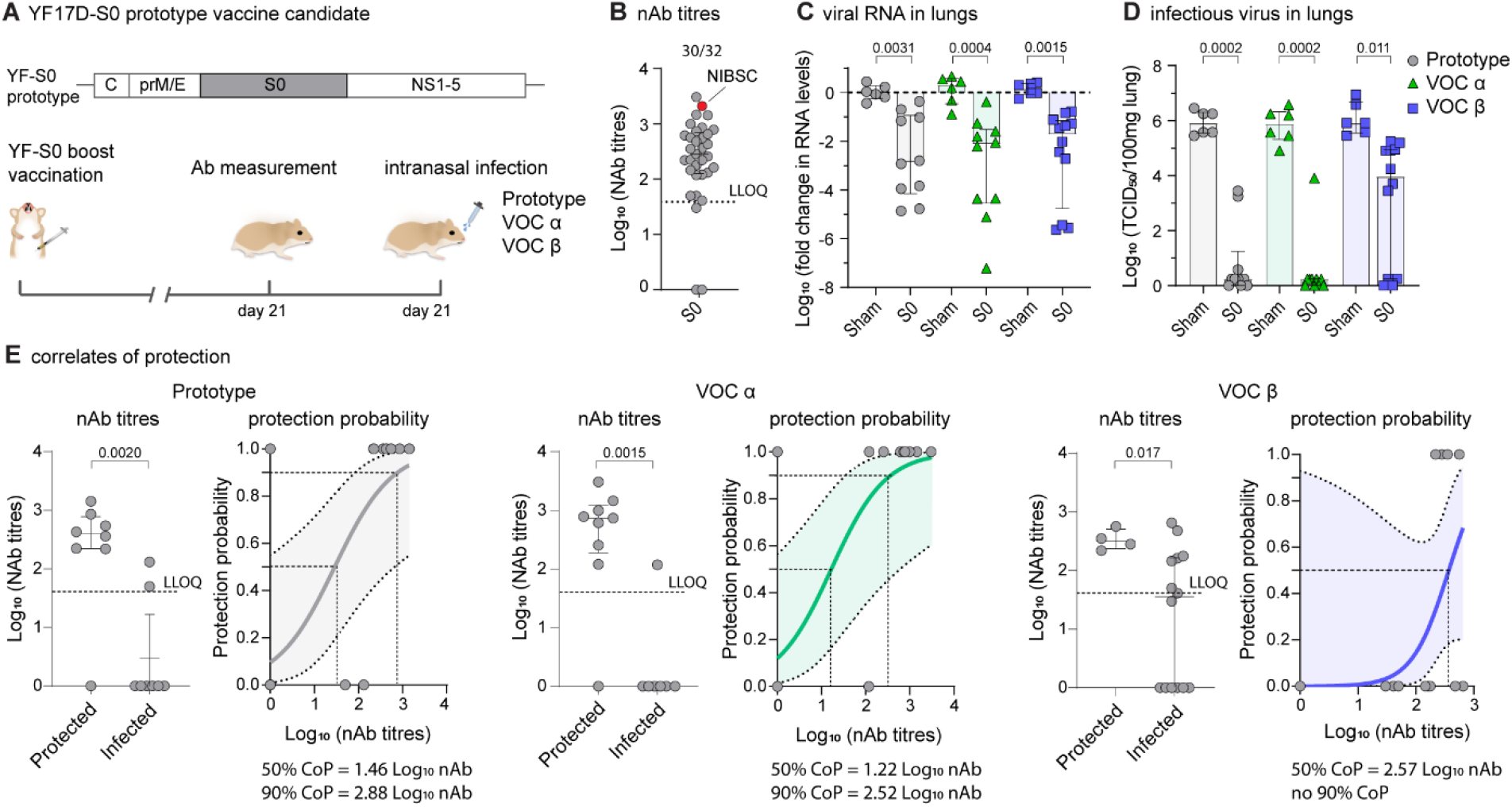
Immunogenicity and protective efficacy of first-generation Spike vaccine against prototype SARS-CoV-2 and VOCs Alpha and Beta. **A**, Vaccination scheme with prototypic YF17D-based vaccine candidate YF-S0 (S0). Syrian hamsters were immunized twice intraperitoneally with 10^4^ PFU of S0 on day 0 and 7 and inoculated intranasally on day 21 with 10^3^ median tissue-culture infectious dose (TCID_50_) of either prototype SARS-CoV-2 (*grey circles*), VOC Alpha (*green triangles*) or VOC Beta (*blue squares*). **B**, nAb titers against prototypic spike (D614G) pseudotyped virus on day 21 after vaccination. Red datapoint indicates the NIBSC 20/130 human reference sample included as benchmark. **C, D**, Viral loads in hamster lungs four days after infection quantified by quantitative RT-PCR (**C**) and virus titration (**D**). **E**, correlates of protection against prototype SARS-CoV-2, VOC Alpha and VOC Beta. Logistic regression model to calculate nAb titers correlating with 50% and 90% probability for protection. ‘*Protected’* was defined by a viral load <10^2^ TCID_50_/100mg lung tissue and ‘*infected’* by a viral load >10^2^ TCID_50_/100mg lung tissue (van der Lubben et al., 2021). Shaded areas indicate 95% CI. LLOQ is lower limit of quantification. Error bars denote median ± IQR. Differences between groups were analyzed using non-parametric Kruskal Wallis test uncorrected for ties.

Logistic regression used to define immune correlates of protection (32) confirmed that comparable nAb levels were required for protection against prototypic SARS-CoV-2 (1.5 log_10_ for 50% and 2.9 log_10_ for 90% protection) and VOC Alpha (1.2 log_10_ for 50% and 2.5 log_10_ for 90% protection) (**Fig. 1E**). Intriguingly, for VOC Beta a markedly (up to 25x) higher nAb threshold (2.6 log_10_) was required for 50% protection. Importantly, no 90% protective nAb threshold could be defined anymore for VOC Beta infection, considering the high number of S0-vaccinated animals with viral breakthrough (>10^2^ TCID_50_/100mg lung tissue) (32). Overall, these data suggest that first-generation vaccines employing prototypic spike as antigen may generally suffer from a markedly reduced efficacy against emerging SARS-CoV-2 variants, such as VOC Beta.

### Updated spike antigen offers complete protection against full range of VOCs Alpha, Beta, Gamma and Delta

Although prototype S0 showed induction of high titres of nAb against prototypic SARS-CoV-2 (**Fig. 1B**) and protective immunity against prototypic SARS-CoV-2 and VOC Alpha (**Fig. 1C-E**), the prototypic spike antigen failed to induce consistent nAb responses against remaining VOCs (**Fig. 2A**). Most importantly, YF-S0 vaccination resulted only in poor seroconversion and low nAb titres against VOC Beta (seroconversion rate 15/32; GMT 1.0 log_10_, 95% CI of 0.6-1.3;) and Gamma (19/32; GMT 1.3 log_10_, 95% CI 0.9-1.8). Intriguingly, human convalescent plasma used as benchmark (WHO standard NIBSC 20/130) originating from 2020 prior to the surge of VOC (**Fig. 2A-B**) showed a similar loss of activity against VOC Beta, in line with what we observed in our hamster sera (**Fig. 2A**).

**Figure 2.**
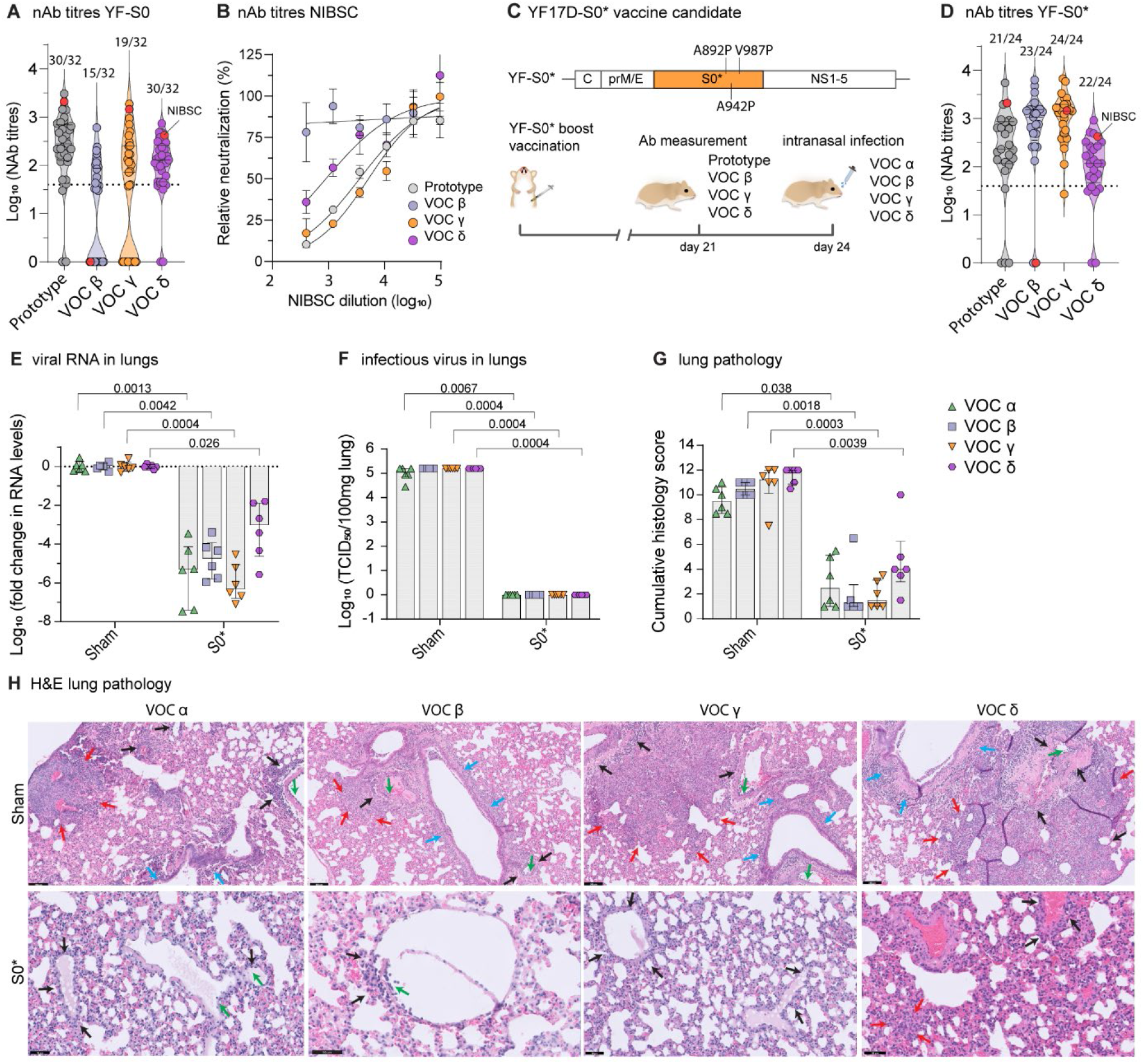
A vaccine based on updated Spike antigen S0* offers complete protection against four VOCs. **A**, nAb titers against prototypic (*grey*), VOC Beta (*blue*), VOC Gamma (*orange*) and VOC Delta (*purple*) spike pseudotyped virus on day 21 after vaccination with prototype YF-S0. Red datapoint indicates the NIBSC 20/130 human reference sample (see Fig. 1B). **B**, Neutralization curves for NIBSC 20/130 human reference sample against same set of pseudotyped viruses. **C**, Schematic of the updated YF-S0* (S0*) vaccine candidate based on VOC Gamma, plus three extra stabilizing proline residues. Vaccination scheme with YF-S0*. Syrian hamsters were immunized twice intraperitoneally with 10^4^ PFU of S0* on day 0 and 7 and inoculated intranasally on day 24 with 10^3^ median tissue-culture infectious dose (TCID_50_) of either VOC Alpha (*green*), VOC Beta (*blue*), VOC Gamma (*orange*) and VOC Delta (*purple*). **D**, nAb titers against prototypic, VOC Beta, VOC Gamma and VOC Delta spike pseudotyped virus on day 21 after vaccination with YF-S0*. Red datapoint indicates the NIBSC 20/130 human reference sample. **E, F**, Viral loads in hamster lungs four days after infection quantified by quantitative PCR with reverse transcription (RT–qPCR) (**E**) and virus titration (**G**). **G**, cumulative lung pathology scores from H&E-stained slides of lungs for signs of damage. **H**, Representative H&E-stained images of sham-or S0*-vaccinated hamster lungs after challenge. Perivascular inflammation (*black arrows*) with focal endothelialitis (*green arrows*); peri-bronchial inflammation (*blue arrows*); patches of bronchopneumonia (*red arrows*). Error bars denote median ± IQR. Differences between groups were analyzed using non-parametric Kruskal Wallis test uncorrected for ties.

It is not clear if the full spectrum of antigenic variability of current VOCs and emerging variants can be covered by a COVID-19 vaccine that is based on a single antigen (33, 34). In an attempt to generate a more universal SARS-CoV-2 vaccine (YF-S0*, S0*), we adapted the spike sequence in our original YF-S0 construct to include the full amino acid spectrum from VOC Gamma, plus three extra proline residues (A892P, A942P and V987P) to stabilize spike in a conformation favorable for immunogenicity (29, 35) (**Fig. 2C**). YF-S0* proved to be highly immunogenic against prototypic SARS-CoV-2, with nAb levels reaching GMT of 2.2 log_10_ (95% CI 1.8-2.6) and a seroconversion rate of 21/24 (**Fig. 2D**), comparable to original YF-S0 (GMT 2.3 log_10_, 95% CI 2.0-2.6; 30/32 seroconversion rate) (**Fig. 2A**). Also, for both constructs, seroconversion rates and nAb levels against VOC Delta were equally high (YF-S0: 30/32; GMT 2.0 log_10_, 95% CI 1.7-2.2; YF-S0*: 22/24; GMT 2.0 log_10_, 95% CI 1.6-2.3). Notably, for YF-S0*, nAb levels and seroconversion rates against VOC Beta (GMT 2.9 log_10_, 95% CI 2.6-3.2; seroconversion rate 23/24) and Gamma (GMT 3.0 log_10_, 95% CI 2.8-3.2; seroconversion rate 24/24) were markedly increased (by 50 to 80-fold for GMT; 1.7 to 2-times more frequent seroconversion) (**Fig. 2A, D**). S0*-vaccinated animals were subsequently challenged with each 10^3^ TCID_50_ of either of the four VOCs Alpha, Beta, Gamma or Delta, and sacrificed 4 dpi for assessment of viral loads in the lung (**Fig. 2E, F**) and associated lung pathology (**Fig. 2G, H**). In S0*-vaccinated hamsters, viral RNA loads were uniformly reduced compared to matched sham controls by ∼3 (VoC Delta) up to ∼6 log_10_ (VoC Gamma) depending on the respective challenge virus under study (**Fig. 2E**). Importantly, no infectious virus could be detected anymore (∼6 log_10_ reduction) in any of the animals vaccinated with S0*, irrespective of which VOC they had been exposed to (**Fig. 2F**), confirming 100% VE conferred by S0* against all four VOCs.

Protection from infection also translated in a markedly reduced pathology (**Fig. 2G, H**). Non-vaccinated sham animals developed characteristic signs of bronchopneumonia with perivascular and peribronchial infiltrations, edema and consolidation of lung tissues (28, 36). In contrast, lungs of S0*-vaccinated hamsters remained markedly less affected with a clear reduction in overall histological scores, irrespectively of the VOC used (**Fig. 2G, H**). In conclusion, second-generation YF-S0* expressing an updated S0* antigen induced consistently high levels of broadly neutralizing antibodies (**Fig. 2D**) which translated into efficient protection from lower respiratory tract infection and COVID-19-like pathology by a large spectrum of VOCs (**Fig. 2G, H**). VE of S0* covered VOCs Beta and Gamma, i.e. variants harbouring key mutations K417N/T and E484K escaping original spike-specific nAb activity (**Fig. 2B**), and may therefore also offer better protection against other emerging variants such as VOI Mu (E484K), or more recently VOC Omicron with similar signatures.

### Blocking of VOC Delta transmission by single dose vaccination

VOC Delta is characterized by a particular efficient human-to-human transmission (37). An added benefit of vaccination at the population level would hence be an efficient reduction in viral shedding and transmission by vaccinated people (38), ideally from single-dose vaccination. For experimental assessment, two groups of hamsters (N=6 each) were either vaccinated once with 10^4^ PFU of S0* or sham (Sanchez-Felipe et al., 2021), and were intranasally infected three weeks later with a high dose comprising 10^5^ TCID_50_ of VOC Delta to serve as index (donor) animals for direct contact transmission (**Fig. 3A**). At 2 dpi, i.e. at onset of increasing viral loads and shedding (39, 40), index animals were each co-housed with one non-vaccinated sentinel for two consecutive days. At 4 dpi, index hamsters were sacrificed, and lungs were assessed for viral RNA, infectious virus and histopathology. Sentinels were sacrificed another two days later and analyzed accordingly.

**Figure 3.**
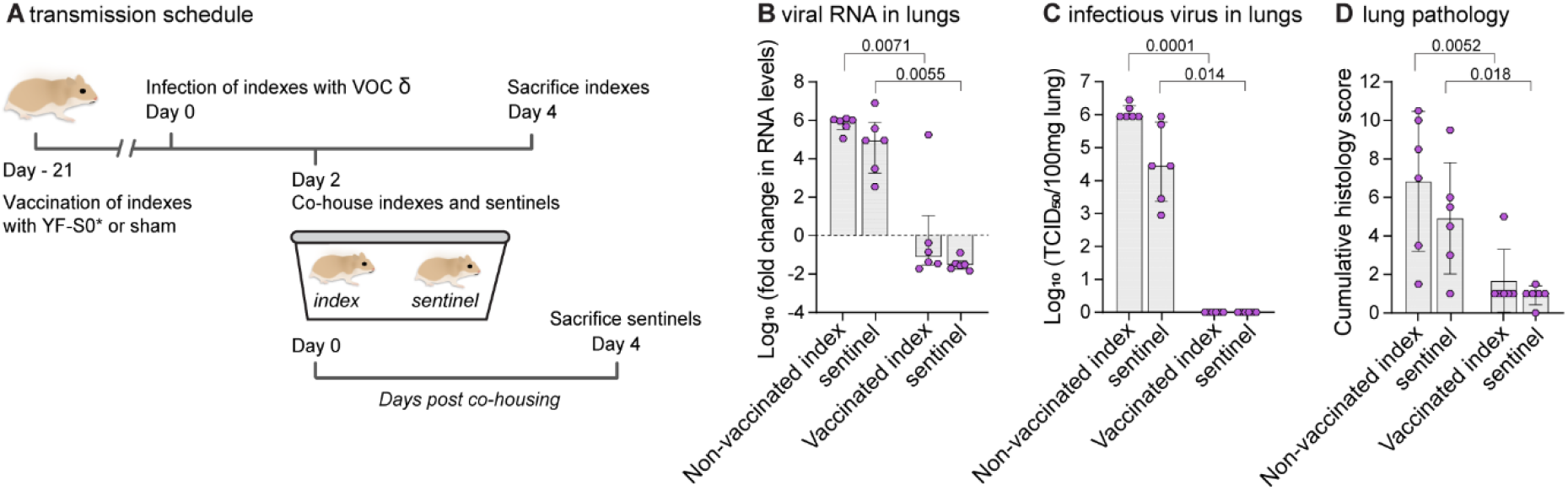
A vaccine based on updated Spike antigen S* completely prevents transmission of VOC Delta. Effect of YF-S0* vaccination on viral transmission to non-vaccinated contact hamsters. Index hamsters were either sham-vaccinated or vaccinated with a single dose of 10^4^ PFU of YF-S0* and infected intranasally on day 21 with 10^5^ TCID_50_ of VOC Delta. Two days after infection, index animals were paired and co-housed with each one naÏve sentinel. Index and sentinel animals were sacrificed each 4 days after infection or exposure, respectively. **B, C**, Viral loads in hamster lungs four days after infection quantified by quantitative RT-qPCR (**B**) and virus titration (**C**). **D**, Cumulative lung pathology scores from H&E-stained slides of lungs for signs of damage. Error bars denote median ± IQR. Differences between groups were analyzed using non-parametric Kruskal Wallis test uncorrected for ties.

As expected from previous experiments, viral loads in S0*-vaccinated index animals were much lower than in non-vaccinated index animals, or than in sentinels that had been in close contact with non-vaccinated donors (**Fig. 3B, C**). Importantly, only very low levels of viral RNA and no infectious virus was observed in non-vaccinated sentinels that had been co-housed with S0*-vaccinated donors. Also, lung pathology was reduced significantly in vaccinated index and co-housed sentinels as compared to sham vaccinated index and respective co-housed sentinels (**Fig. 3D**). To our knowledge, this constitutes the first experimental evidence for full protection from SARS-CoV-2 transmission by any vaccine. The block conferred by S0* appears to be more complete than that observed in humans by current vaccines (41).

### Increased potency of new vaccine candidate S0* against Omicron variant

Immunogenicity of S0 and S0* was tested in parallel against VOC Omicron. Sera from hamsters vaccinated with updated S0* resulted in 60% (19/32) seroconversion to nAbs as compared to 4% (2/44) in those vaccinated with the prototypic S0 construct. Both vaccine candidates S0 and S0* were equally active against prototype (Kruskal-Wallis multiple groups; *p*= 0.79). Quantitatively, S0* resulted in a marked ∼15-fold increase (p<0.0001) in nAbs with activity against Omicron to log_10_ GMT of 1.2 (IQR 0-2.06), compared to S0 which resulted in hardly any nAb against Omicron (log_10_ GMT of 0.09 log_10_ IQR 0-0.09), confirming the substantial gain in immunogenicity for S0* (**Fig. 4A**).

**Figure 4.**
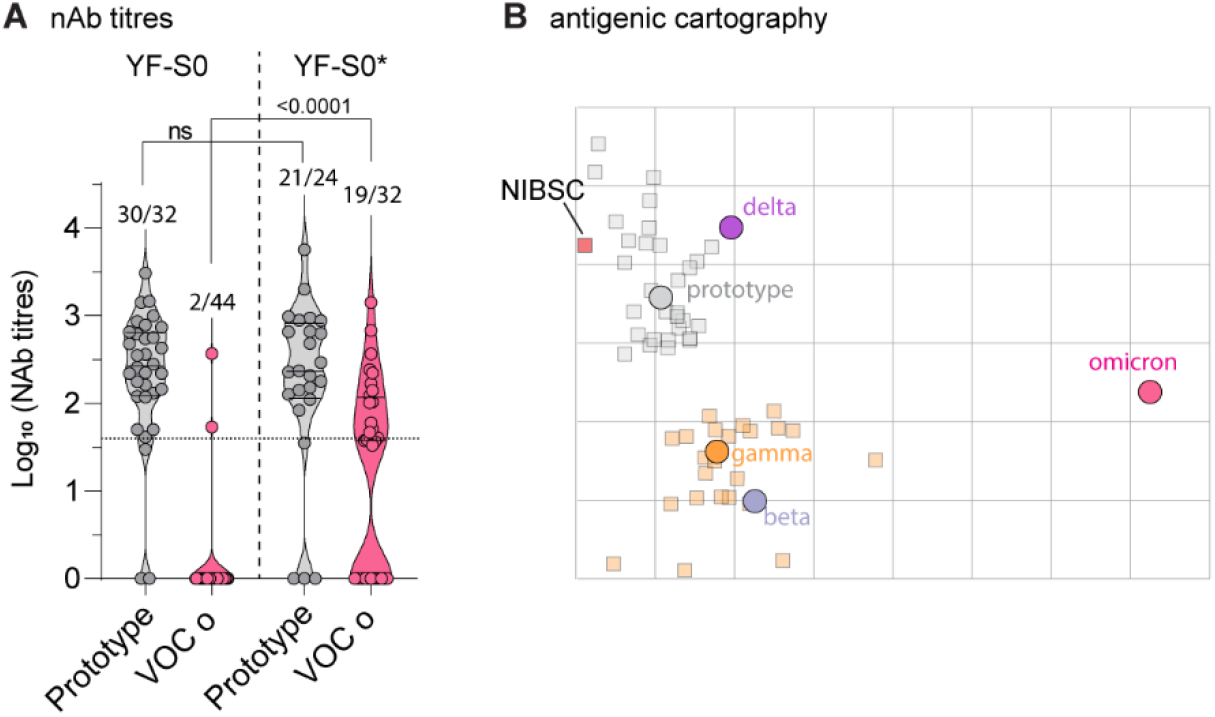
Increased potency of new vaccine candidate S0* against VOC Omicron. **A**, nAb titers against prototypic (*grey*) and omicron (*pink*) spike pseudotyped virus on day 21 after vaccination with prototype YF-S0 or YF-S0*. Error bars denote median ± IQR. Differences between groups were analyzed using non-parametric Kruskal Wallis test uncorrected for ties. **B**, Antigenic cartography. Cross-reactivity of sera raised by original S0 (*grey squares*) and updated S0* (*orange squares*) vaccine antigen against five different SARS-COV-2 variants (circles: prototype, *grey*; VOC Beta, *blue*; Gamma, *orange*; Delta, *purple*; Omicron, *pink*) plotted on a two-dimensional map (Smith et al., 2004). NIBSC, human reference serum pool (NIBSC 20/13).

This change in cross-reactivity of the sera raised by the original (YF-S0) and updated (YF-S0*) vaccine antigen against five different virus variants (prototype; VOCs Beta, Gamma, Delta and Omicron) was further studied using antigenic cartography (42) (**Fig. 4B)**. VOC Alpha was not considered since it did not differ from the prototype virus, neither regarding VE of S0 nor nAb titre as correlate protection (**Fig. 1**). Specifically, we constructed a two-dimensional projection that geometrically maps median serum neutralization titres (SNT_50_) between sera and respective antigens as antigenic distances. This revealed a pattern of antigenic diversification between prototype virus on the one hand and VOCs Beta and Gamma on the other hand, with VOC Delta being mapped closer to the prototype virus as compared to Beta and Gamma. This is consistent with recently described patterns of convergent evolution in spike for VOCs Beta and Gamma, and Delta climbing a different fitness peak (43). VOC Omicron appeared by far more distant from any other strain, in line with recent larger scale antigenic analysis (44).

In line with the visual pattern of clustering, antigenic distances for S0* sera were significantly larger to prototype and VOC Delta than to Beta and Gamma (t-test; p<0.001). Intriguingly, however, this obvious antigenic drift did not reduce the overall higher potency of S0*, which included equally strong humoral responses to prototypic spike and VOC Delta (**Fig. 2A, D**). Likewise, despite being antigenically still largely divergent from other VOC (mean range: 0.8-3.3 units), antigenic distances regarding VOC Omicron were significantly shorter (t-test; p<0.0001) for S0* sera compared to S0 sera (respectively mean ± SD; 5.5 ± 0.7 and 6.4 5 ± 0.5), in further support for the marked gain in Omicron-specific humoral immunity by S0* (**Fig. 4A**). Overall, these findings support the general observation that vaccines employing prototypic spike as antigen are losing serological coverage, in particular towards those VOCs (Beta, Gamma and Omicron) linked to escape from antibody-mediated immunity. We further demonstrate how vaccine potency and induction of cross-reacting nAb can be markedly enhanced by alternative spike antigen choice and design.

## Discussion

Little experimental support exists on how well current first-generation vaccines protect against the full spectrum of VOCs. While likely protecting from severe COVID-19 caused by any SARS-CoV-2 strain, a clear drop in VE was observed during clinical trials conducted in regions with high circulation of VOC Beta as paradigm of an E484K Spike variant and others known to escape nAb recognition (45). Experimentally, such a drop in protective immunity is confirmed by higher viral loads in macaques vaccinated with an Adenovirus-vectored prototype spike antigen (Ad26.COV2.S) and challenged with VOC Beta (46). Likewise, in the more stringent hamster model, immunity acquired during previous SARS-CoV-2 (prototype) infection, or by Ad26.COV2.S vaccination, led only to partial suppression of heterologous VOC Beta replication (47). In the latter case, replicative viral RNA was still detectable two weeks after challenge (< 2 log_10_ reduction compared to sham), which is completely in line with the observed failure of prototypic YF-S0 to confer full protection against VOC Beta (**Fig. 1F-H**). By contrast, viral replication was reduced to undetectable levels for all four VOCs by YF-S0* vaccination using an updated spike antigen (**Fig. 2F**). Finally, S0* blocked transmission of VOC Delta, for which a single dose vaccination was sufficient (**Fig. 3**). The stringent hamster model is generally well suited to assess both aspects of preclinical VE, individual protection and transmission (24-26, 48). An obvious shortcoming of our current study is the limited infectivity of VOC Omicron in hamsters (30), and that VE of YF-S0*against Omicron can hence not directly be assessed in this gold standard model. However, considering (*i*) that VOC Omicron is poorly, if at all, covered by current vaccines (50) and (*ii*) nAbs are strongly correlated with VE, the gain in Omicron-specific nAb achieved by YF-S0* vaccination is remarkable. Successful testing in complementary animal models (e.g. human ACE2-transgenic hamsters (50) will warrant further development of YF-S0* as a second-generation COVID-19 vaccine candidate with broader coverage of relevant virus strains.

In more general terms, our findings strongly suggest that first-generation COVID-19 vaccines will need to be adapted to keep up with the evolution of variants driving the ongoing global SARS-CoV-2 pandemic as a result of their critical combinations of driver mutations responsible for both nAb escape and enhanced transmission.

## Methods

### Viruses and animals

All virus-related work was conducted in the high-containment BSL3 facilities of the KU Leuven Rega Institute (3CAPS) under licenses AMV 30112018 SBB 219 2018 0892 and AMV 23102017 SBB 219 2017 0589 according to institutional guidelines. All SARS-CoV-2 strains used throughout this study were isolated in house (University Hospital Gasthuisberg, Leuven) and characterized by direct sequencing using a MinION as described before (36). Strains representing prototypic SARS-CoV-2 (Wuhan; EPI_ISL_407976) (36), VOCs Alpha (B.1.117; EPI_ISL_791333) and Beta (B.1.351; EPI_ISL_896474) have been described (28). Strains representing VOCs Gamma (P.1; EPI_ISL_1091366) and Delta (B.1.617.2; EPI_ISL_2425097) were local Belgian isolates from March and April 2021, respectively.

All virus stocks were grown on Vero E6 cells and used for experimental infections at low *in vitro* passage (P) number, P3 for prototype and P2 for all four VOCs. Absence of furin cleavage site mutations was confirmed by deep sequencing. Median tissue culture infectious doses (TCID_50_) were defined by titration as described (28, 36) using Vero E6 cells as substrate, except for VOC Delta, for which A549 cells were used for a more pronounced virus induced cytopathic effect (CPE). Housing and experimental infections of hamsters have been described (1, 36, 39) and conducted under supervision of the ethical committee of KU Leuven (license P050/2020 and P055/2021). In brief, 6 to 8 weeks old female Syrian hamsters (*Mesocricetus auratus*) were sourced from Janvier Laboratories and kept per two in individually ventilated isolator cages. Animals were anesthetized with ketamine/xylazine/atropine and intranasally infected with 50 μL of virus stock (25 μL in each nostril) containing either 10^3^ or 10^5^ TCID_50_ as specified in the text and euthanized 4 days post infection (dpi) for sampling of the lungs and further analysis. Animals were monitored daily for signs of disease (lethargy, heavy breathing, or ruffled fur).

### Vaccine Candidate

The general methodology for the design and construction of a first YF17D-based SARS-CoV-2 vaccine candidate (YF-S0) has been described (1). Several mutations were introduced into original YF-S0 to generate second-generation vaccine candidate YF-S0*. The first series of mutations is based on the spike sequence of VOC Gamma: L18F, T20N, P26S, D138Y, R190S, K417T, E484K, N501Y, D614G, H655Y, T1027I, V1176F. A second series of mutations is based on a locked spike variant, stabilizing the protein in a more immunogenic prefusion confirmation: A892P, A942P, V987P (29).

### Production of spike-pseudotyped virus and serum neutralization test (SNT)

Virus-neutralizing antibodies (nAb) were determined using a set of VSV spike-pseudotype viruses essentially as described (1). For this purpose, five different pseudotypes were generated using expression plasmids of respective spike variants: for prototype B.1/D614G as before (1) or sourced from Invivogen for VOCs Beta (Cat. No. plv-spike-v3), Gamma (Cat. No. plv-spike-v5) and Delta (Cat. No. plv-spike-v8). The VOC Omicron spike expression construct was assembled from six custom synthetized gDNA fragments (IDT, Leuven) and cloned into pCAG2 plasmid backbone as before (1). Briefly, depending on the plasmid background, BHK-21J cells (variant B.1/D614G and Omicron) or HEK-293T cells (Beta, Gamma and Delta) were transfected with the respective SARS-CoV-2 spike protein expression plasmids, and one day later infected with GFP-encoding VSVΔG backbone virus (51). Two hours later, the medium was replaced by medium containing anti-VSV-G antibody (I1-hybridoma, ATCC CRL-2700) to neutralize residual VSV-G input. After 26 hours incubation at 32°C, the supernatants were harvested. To quantify nAb, serial dilutions of serum samples were incubated for 1 hour at 37 °C with an equal volume of S-pseudotyped VSV particles and inoculated on Vero E6 cells for 19 hours.

The resulting number of GFP expressing cells was quantified on a Cell Insight CX5/7 High Content Screening platform (Thermo Fischer Scientific) with Thermo Fisher Scientific HCS Studio (v.6.6.0) software. Median serum neutralization titres (SNT_50_) were determined by curve fitting in Graphpad Prism after normalization to virus (100%) and cell controls (0%) (inhibitor *vs*. response, variable slope, four parameters model with top and bottom constraints of 100% and 0%, respectively).The human reference sample NIBSC 20/130 was obtained from the National Institute for Biological Standards and Control, UK.

### Antigenic cartography

We used the antigenic cartography approach developed for influenza hemagglutination inhibition assay data to study the antigenic characteristics of the SARS-CoV-2 Spikes (42). This approach transforms SNT_50_ data to a matrix of immunological distances. Immunological distance *d*_*ij*_ is defined as *d*_*ij*_ = *s*_*j*_ – *H*_*ij*_, where *H*_*ij*_ is the log_2_ titre of virus *i* against serum *j* and *s*_*j*_ is the maximum observed titre to the antiserum from any antigen (*s*_*j*_ = max(*H*_*1j*_, …, *H*_*nj*_)). Subsequently, a multidimensional scaling algorithm was used to position points representing antisera and antigens in a two-dimensional space such that their distances best fit their respective immunological distances. Even though distances are measured between sera raised by vaccination using specific Spike antigens (and the NIBSC serum) and antigens, such an antigenic map also provides estimates of antigenic distances between the antigens themselves.

### Vaccination and challenge

COVID-19 vaccine candidate YF-S0 (1) was used to vaccinate hamsters at day 0 and day 7 (N=32) with a dose of 10^4^ PFU via the intraperitoneal route and control animals (N=18) were dosed with MEM (Modified Earl’s Minimal) medium containing 2% bovine serum as sham controls. Blood was drawn at day 21 for serological analysis and infection was done on the same day with prototype (N=10 vaccinated; N=6 sham), VOC Alpha (N=10 vaccinated; and N=6 sham), and VOC Beta (N=12 vaccinated; N=6 sham) with the inoculum of 10^3^ TCID_50_ intranasally. Protective nAb levels were calculated using logistic regression analysis in GraphPad Prism (version 9) as described (32)

Similarly, hamsters were vaccinated twice with 10^4^ YF-S0* (N=24) or sham (N=16) at day 0 and day 7. Blood was collected at day 21 to analyze nAbs in serum, and animals were infected on day 24 with different variants, including VOCs Alpha, Beta, Gamma and Delta with the inoculum of 10^3^ TCID50 intranasally (N=6 vaccinated and N=4 sham vaccinated infected against each variant). Lungs were collected for analysis of viral RNA, infectious virus and for histopathological examination as described in (1). Resulting vaccine efficacy (VE) was calculated as [1 – (number of vaccinated animals with detectable virus) / (number of all infected animals)] x 100% per group of hamsters infected with the same virus strain, whereby a lung viral load >10^2^ TCID_50_/100mg was set as cutoff for infection (32).

To assess VE of S0* against Omicron, hamsters were vaccinated twice at day 0 and day 7 with 10^4^ PFU YF-S0* (N=12) and YF-S0 (N=12). Blood was collected on day 28 to analyze nAbs against Omicron. In parallel, samples from previous experiments (day 21) were also tested for nAbs against Omicron.

### Viral load and viral RNA quantification

Virus loads were determined by titration and RT-qPCR from lung homogenates was performed exactly as previously described in detail (1, 36, 39).

### Histopathology

For histological examination, the lungs were fixed overnight in 4% formaldehyde, embedded in paraffin and tissue sections (5 μm) after staining with H&E scored blindly for lung damage (cumulative score of 1 to 3 each for congestion, intra-alveolar hemorrhage, apoptotic bodies in bronchial epithelium, necrotizing bronchiolitis, perivascular edema, bronchopneumonia, perivascular inflammation, peribronchial inflammation, and vasculitis) as previously established (28, 36)

### Blocking of viral transmission

Hamsters (N=6) were vaccinated with 10^4^ PFU of vaccine once, were bled at day 21 and infected with VOC Delta with 1×10^5^ TCID50, intranasally. Another group of non-vaccinated hamsters (N=6) were also infected. Two days post infection index animals were co-housed with sentinels for two days and separated after two days of exposure. All the index animals were euthanized on day four post infection and sentinels were sacrificed after 4 days of exposure. Lungs were analyzed for viral RNA and infectious virus and subjected to histopathology.

### Statistical analysis

All statistical analyses were performed using GraphPad Prism 9 software (GraphPad, San Diego, CA, USA). Results are presented as GM± IQR or medians ± IQR as indicated. Data were analyzed using uncorrected Kruskal-Wallis test and considered statistically significant at p-values ≤0.05.

## Acknowledgements

We are grateful to Prof. Michael A. Whitt (University of Tennessee Health Science Center) for generously sharing of plasmids for rescue of VSVΔG, and Dr. Maya Imbrechts and Dr. Nick Geukens (PharmAbs, KU Leuven) for help with hybridoma culture. We thank Carolien De Keyzer and Lindsey Bervoets (Rega) for excellent technical assistance with animal experimentation as well as the staff of the Rega animalium for strong support. We thank Jasper Rymenants, Tina van Buyten, Dagmar Buyst, Thibault Francken, Niels Cremers and Stijn Hendrickx for steady and timely help with analysing of tissue samples and virus titrations. We finally thank Jasmine Paulissen, Catherina Coun, Céline Sablon, Jolien Timmermans and Nathalie Thys (TPVC) for diligent serology assessment, vaccine stock generation and skilled generation of plasmid constructs.

Current work was supported by the Flemish Research Foundation (FWO) emergency COVID-19 fund (G0G4820N) and the FWO Excellence of Science (EOS) program (No. 30981113; VirEOS project), the European Union’s Horizon 2020 research and innovation program (No 101003627; SCORE project and No 733176; RABYD-VAX consortium), the Bill and Melinda Gates Foundation (INV-00636), KU Leuven Internal Funds (C24/17/061) and the KU Leuven/UZ Leuven Covid-19 Fund (COVAX-PREC project) and European Health Emergency Preparedness and Response Authority (HERA). G.B. acknowledges support from the KU Leuven Internal Funds (Grant No. C14/18/094) and the Research Foundation - Flanders (“Fonds voor Wetenschappelijk Onderzoek - Vlaanderen,” G0E1420N, G098321N). P.L. acknowledges funding from the European Research Council under the European Union’s Horizon 2020 research and innovation program (grant agreement no. 725422-ReservoirDOCS) and from the EU grant 874850 MOOD. K.D. acknowledges grant support from KU Leuven Internal Funds (C3/19/057 Lab of Excellence).

## Contributions

S.S. and K.D. conceptualization; S.S. animal experimentation; S.S., T.V., W.K., M.R. and H.J.T. data generation, analysis and curation; S.S., H.J.T. and K.D. original manuscript draft; S.S. and H.J.T. visualization; T.V. and L.S.F. construct design; T.V., W.K. and D.V.L. serological analysis; R.A. and C.S.F. VoC hamster models; B.W. histological analysis; P.L. and G.B. antigenic cartography; L.S.F., V.L., T.V., W.K., and P.M. vaccine stocks and virus isolation; J.N., H.J.T., and K.D. supervision, writing and project administration; J.N. and K.D. funding acquisition. All authors read, edited and approved the final version of the manuscript.

## Notes

### Competing Interest Statement

The authors have declared no competing interest.

### Summary of Updates

Data set on VOC Omicron added comprising nAb responses as elicited by prototype and updated vaccine antigen (presented in new Figure 4). Antigenic cartography adapted accordingly to including VOC Omicron data. Title, abstract and main conclusions have been revised and adapted accordingly. New author added.

